# A Variational Modeling Framework for Population Genetic Dynamics

**DOI:** 10.64898/2026.09.17.750900

**Authors:** Cai Li

## Abstract

Population genetic dynamics are shaped by multiple evolutionary processes, including mutation, recombination, selection, and genetic drift. With the rapid growth of genomic data, modeling multilocus evolutionary dynamics and the resulting patterns of genetic variation has become increasingly important. Jointly modeling multiple evolutionary processes across multiple loci remains challenging, motivating the development of more flexible and extensible frameworks. Here, we develop a variational framework for population genetic dynamics based on generalized gradient-flow theory, drawing on nonequilibrium thermodynamics. In our model, mutation, recombination, and selection are represented as modular variational components, connected through a gamete–individual two-state system with active coupling. Mutation and recombination share a gamete-state free energy but use distinct dissipation potentials, while selection uses the established Shahshahani gradient structure at the individual stage. Our model represents multilocus genetic variation directly in the full haplotype-frequency space, recovers classical mutation, recombination, and selection dynamics in the corresponding limits, and provides a phenomenological stochastic extension for finite-population genetic drift. Numerical comparisons with SLiM forward simulations support consistency with allele-frequency, haplotype-frequency, and linkage-disequilibrium dynamics in the tested systems. This variational formulation provides a new perspective on modeling evolutionary dynamics and has potential for extension to more complex multilocus systems and applications in population genomics analysis.

## 1 Introduction

### 1.1 Motivation and challenges in modeling population genetic dynamics

Over the past century, population genetics has built a rich body of theories and methods that laid the foundation for understanding intraspecific genetic variation and its evolutionary history. However, as the scale and complexity of genomic data grow rapidly, many classical theories still rely on specific simplifying assumptions and mathematical approximations whose generalizability remains limited, restricting their ability to capture the full range of evolutionary information encoded in large-scale genomic data [1]. These limitations highlight the need for theoretical frameworks and approximation methods that can accommodate new genomic data and research questions [2].

Population genetic dynamics involve evolutionary processes with different dynamical structures, including mutation, recombination, selection, and drift [3]; their combined effects increase model complexity, and how to describe multiple evolutionary processes in a unified way remains an important theoretical challenge [4]. For multilocus systems, the state space grows rapidly with the number of loci, so a complete description of haplotype frequencies and inter-locus associations faces significant dimensionality and computational challenges [5].

Classical multilocus theory addresses these dynamics through recursion equations for selection, recombination, and genetic associations [6], with extensions to mutation, migration, and diverse inheritance systems [7], but exact representations remain high-dimensional and quasi-linkage equilibrium (QLE) approximations require selection to be weak relative to recombination. Analytical approaches include exact solutions for independent mutation, single-crossover recombination, and additive selection [8], and probabilistic representations using random partitions of loci to describe recombination, with labels incorporating selection and mutation [9]. Poulton et al. [10] obtained exact stationary distributions for a finite-population selection–mutation– recombination–drift model through Gibbs sampling, but this tractability relies on n-parent recombination that enforces linkage equilibrium among offspring rather than standard two-parent recombination.

Over the past two decades, several complementary computational approaches have substantially expanded the range of multilocus population-genetic models that can be analyzed. Coalescent and ancestral recombination graph methods represent correlated genealogies generated by recombination and increasingly enable genome-scale reconstruction and inference of demographic history and selection [11,12]. Forward-time simulation frameworks can explicitly incorporate mutation, recombination, selection, drift, and complex demography, while tree-sequence recording has greatly improved their scalability to large genomic regions [13,14]. Complementary diffusion- and moment-based approaches provide tractable descriptions of two-locus dynamics, with methods incorporating recombination, demographic history, drift, and selection [15,16,17]. These approaches address multilocus complexity through genealogical inference, forward simulation, and diffusion- or moment-based modeling. Despite these advances, systematically constructing and extending models that combine multiple evolutionary processes remains challenging. This motivates the search for shared mathematical principles that can guide model development.

### 1.2 Variational methods as a modeling tool for population genetic dynamics

Variational methods have a long history of application in physics and chemistry, where they are used to construct dynamical models from the energy, action, or dissipation potential of a system and to derive the corresponding evolution equations [18]. This variational way of thinking provides a systematic approach to describing dynamical processes through their driving forces and kinetic responses.

In population genetics, early theoretical work established variational and geometric approaches to selection dynamics. Svirezhev and Behera [19,20] formulated selection dynamics in terms of variational or steepest-ascent principles and explored their extension to multilocus systems, while Shahshahani [21] introduced a Riemannian metric that provides a geometric formulation of deterministic selection dynamics. From a statistical-physics perspective, Sella and Hirsh [22] established a close analogy between finite-population evolutionary dynamics and thermodynamic systems, showing that the steady-state distribution of fixed genotypes has a Gibbs-like form and that a free-fitness function characterizes the balance between selection and stochasticity. Rastegar-Sedehi et al. [23] formulated a diploid genotype master equation incorporating selection, mutation, recombination, and reproduction, and analyzed the associated entropy production. Xu and Wang [24] developed a potential landscape and flux framework to characterize the gradient and non-gradient contributions to multilocus evolutionary dynamics. Recently, explicit variational formulations based on generalized gradient-flow theory have also been established for specific evolutionary processes, including recombination as a mechanism that dissipates linkage disequilibrium [25] and the Kimura equation with selection and genetic drift in Wasserstein– Shahshahani geometry [26]. These studies provide variational, geometric, and statistical-mechanical perspectives on population genetic dynamics.

### 1.3 Framework overview and paper organization

Building on this body of work, we adopt variational approaches from nonequilibrium thermodynamics and generalized gradient-flow theory [27] to formulate the deterministic components of population genetic dynamics. The variational formulation specifies the state variables, evolutionary driving forces, and process-specific dissipation potentials, from which the deterministic evolution equations are derived, while genetic drift is incorporated separately through a stochastic extension. Our framework combines three key elements: a gamete–individual two-state system, in which mutation and recombination act on the gamete state and selection acts on the individual state; a modular variational structure, in which different evolutionary processes are represented by process-specific driving forces and dissipation potentials and connected through a gamete–individual coupling structure; and a stochastic extension for genetic drift. This formulation provides a composable mathematical structure for population genetic dynamics, making it amenable to future differentiable implementations and gradient-based computation.

The paper is organized as follows. Section 2 introduces the biological processes and motivates the two-state representation. Section 3 presents the general variational framework, while Section 4 develops the variational formulations of mutation, recombination, and selection. Section 5 integrates these processes into the complete gamete–individual model, and Section 6 introduces the stochastic extension for genetic drift. Section 7 presents numerical experiments, followed by discussion and conclusions in Section 8.

## 2 Evolutionary Processes and the Gamete–Individual Life Cycle

This section introduces the four evolutionary processes considered in the framework and the stages of the life cycle at which they act, motivating the two-state representation used throughout the paper.

### 2.1 Life cycle and biological stages of evolutionary processes

Changes in the genetic composition of a population are mainly driven by four evolutionary processes considered in this framework: mutation, recombination, selection, and genetic drift [3,28]. Population genetic processes can be viewed as a life cycle in which a gamete pool gives rise to a population of individuals, which undergoes selection and produces gametes that form the gamete pool of the next generation. Mutation and recombination are modeled at the gamete-formation stage, whereas selection primarily acts on the survival and reproductive contributions of individuals. Selection can also occur at the gamete stage, for example through meiotic drive or gamete competition, but these effects are not explicitly modeled here.

Kirkpatrick et al. [7] explicitly track gametes, zygotes, adults, and new gametes using contexts that distinguish stage, sex, and parental origin. Their discrete-generation recursions propagate allele frequencies and associations through selection and transmission. We use a simpler continuous-time representation of haploid populations: gamete-level and individual-level haplotype distributions evolve as coupled variables, without separately resolving zygote-to-adult development, sex, or parental origin.

Finite-population sampling fluctuations may arise during both gamete production and the formation of the next generation of individuals. Motivated by these two life-cycle transitions, we introduce Wright– Fisher-type sampling noise in both state variables as a phenomenological stochastic extension (Section 6).

Although the different evolutionary processes jointly change the genetic composition of a population, they do not all act at the same stage of the life cycle (**Fig. 1**).

**Figure 1:**
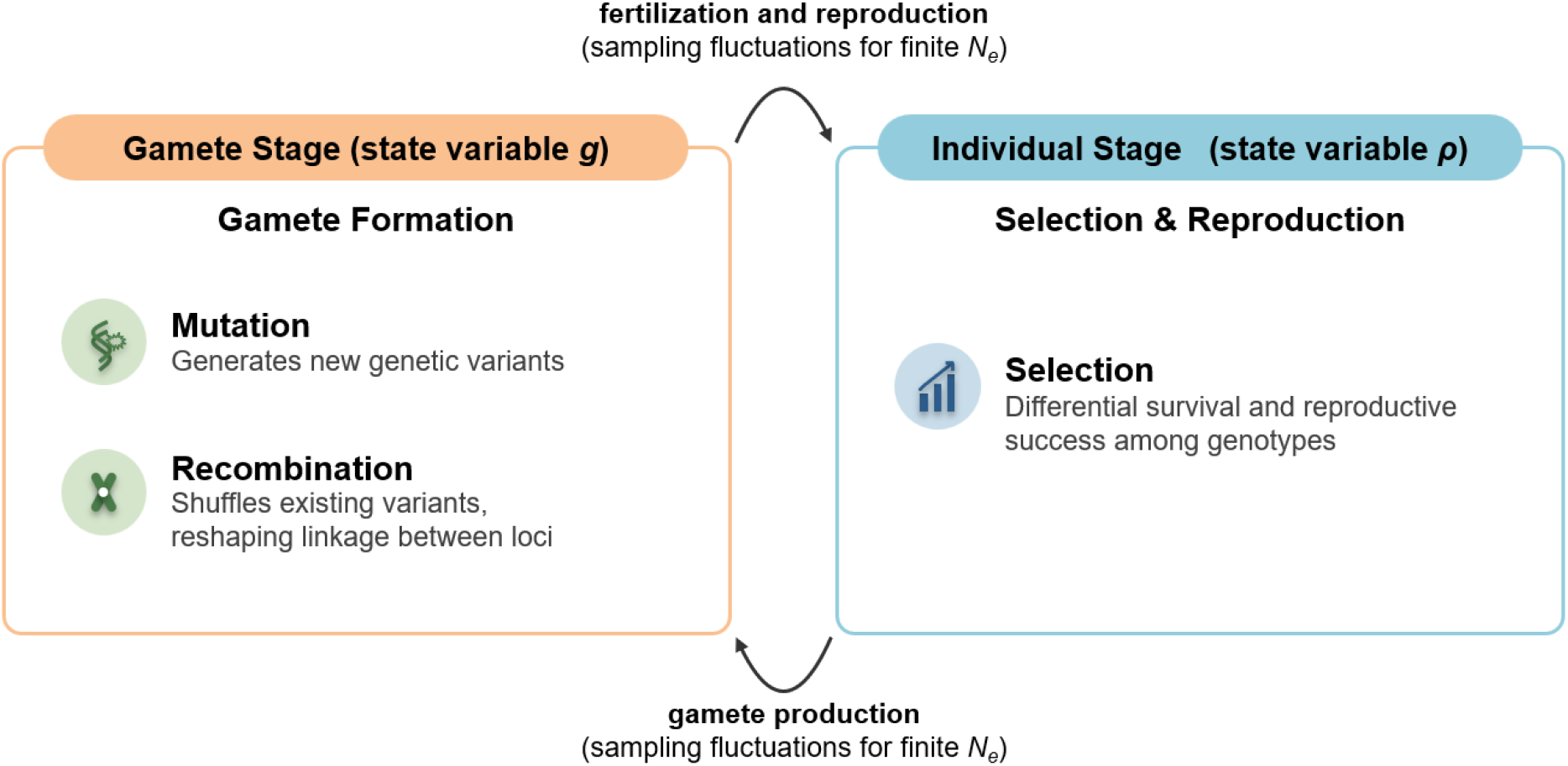
Two stages of the life cycle and associated population genetic processes. In the gamete stage (state variable **g**), mutation generates genetic variation and recombination reshapes linkage disequilibrium. In the individual stage (state variable **ρ**), selection changes the survival and reproductive contributions of haplotypes. Genetic drift is represented phenomenologically by Wright–Fisher-type sampling noise in both state variables, motivated by finite-population sampling at the two life-cycle transitions (Section 6). N_e_, effective population size.

This life-cycle structure motivates two state variables: ***g***, describing the haplotype-frequency distribution in the gamete pool, and ***ρ***, describing the corresponding distribution in the individual population. The two-state representation is not intended to increase mathematical complexity but to reflect the biological separation of life-cycle stages and thus provides the basis for assigning mutation, recombination, selection, and drift to their corresponding dynamical components.

### 2.2 Two-state variable representation

#### 2.2.1 Haplotype space

Consider a haploid population with *L* biallelic loci; denoting the two alleles by 0 and 1, a haplotype is represented as

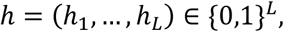

There are therefore 2^*L*^ possible haplotypes, which form the discrete genetic state space for the subsequent dynamics.

#### 2.2.2 Gamete state

The gamete state at time *t* is denoted by

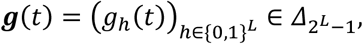

where *g*_*h*_ denotes the frequency of haplotype *h* in the gamete pool and 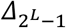 denotes the (2^*L*^ − 1)-dimensional probability simplex:

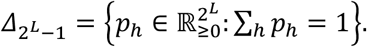

The gamete state describes haplotype frequencies in the gamete pool before the formation of the next generation. Mutation changes the allelic state at specific loci, whereas recombination reshapes linkage disequilibrium among loci; both processes can alter haplotype frequencies.

#### 2.2.3 Individual state

The individual state at time *t* is denoted by

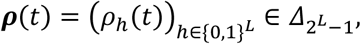

The individual state ***ρ*** is the haplotype-frequency vector in the population of individuals. We focus mainly on haploid populations, so *ρ*_*h*_ denotes the frequency of individuals carrying haplotype *h*. Selection changes the reproductive contributions of different haplotypes and acts primarily on the individual state. Finite-population sampling fluctuations are represented phenomenologically by stochastic terms in both ***g*** and ***ρ*** (Section 6). The extension to diploid populations is given in Section 5.5.

#### 2.2.4 Reproduction map between the two stages

The gamete state ***g*** and the individual state ***ρ*** describe two adjacent stages of the life cycle. Their biological correspondence is specified by a reproduction map:

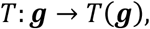

In the haploid setting considered here, the reproduction map preserves haplotype composition, so *T*(***g***) = ***g***. For diploid populations, reproduction generally involves gamete fusion and a mapping from gamete frequencies to genotype distributions (Section 5.5).

Importantly, the reproduction map specifies the expected correspondence between the two stages rather than imposing an instantaneous constraint ***ρ*** = *T*(***g***). In our variational model, ***g*** and ***ρ*** remain distinct dynamical variables, and their discrepancy is regulated by the coupling mechanism introduced in Section 5. In the strong-coupling limit, ***ρ*** approaches *T*(***g***). Thus, the reproduction map defines the biological correspondence, whereas the coupling mechanism establishes this correspondence dynamically.

## 3 Variational Modeling Based on the Energy–Dissipation Principle

This section presents the general variational framework, in which dynamical equations are derived from driving forces and process-specific dissipation potentials. Specific evolutionary processes are treated in Sections 4 and 5, with detailed derivations and proofs provided in the Appendix.

### 3.1 Basic ideas of variational modeling

The variational modeling approach adopted here is based on the energy–dissipation principle of nonequilibrium statistical physics [27]. Here, “variational” means that the evolution equation is derived from an optimization principle over admissible instantaneous rates of change, with the current state held fixed. In this framework, the free energy specifies the thermodynamic driving forces, while dissipation potentials characterize how the system responds to these forces. For a standard passive dissipative process, the instantaneous evolution rate minimizes the sum of the free-energy change rate and the dissipation cost. Thus, the dynamics arise from a balance between energetic driving and kinetic resistance, rather than from minimizing the free energy directly at each instant. By specifying the driving forces and kinetic responses separately, different evolutionary processes can share the same free energy while retaining distinct dissipation potentials, as demonstrated for mutation and recombination in Section 4.1. For systems with active driving, such as the coupled model introduced in Section 5, the total free energy need not be a Lyapunov function of the full dynamics.

Generalized gradient flows provide a mathematical formulation of this variational structure through a state space, an energy functional, and a dissipation potential [27]. Let ***x*** denote the system state and ℱ(***x***) the free energy. Following the terminology of nonequilibrium thermodynamics, we define the thermodynamic force as the negative variational derivative of the free energy,

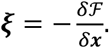

For a dual dissipation potential *ψ*^*^ that is differentiable in the force variable, as in the constructions used here at interior states, the dynamical equation is

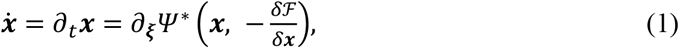

where ∂_*ξ*_*ψ*^*^ denotes the derivative with respect to *ξ*. For a nonsmooth convex potential, this relation is replaced by a subdifferential inclusion.

For a single dynamical process *i*, we denote its dual dissipation potential by 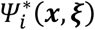. When multiple processes are represented within the same state variable and share the same driving force, their dissipation potentials can be combined as

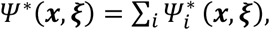

giving

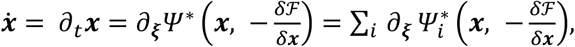

This formulation allows different evolutionary processes to be incorporated as distinct dissipative components within a common variational framework. In this paper, mutation, recombination, and selection are represented by process-specific dissipation potentials. The underlying mathematical theory of generalized gradient flows is provided in [27,29,30].

### 3.2 Primal and dual dissipation potentials

Within the energy–dissipation principle, the dissipation structure can be expressed through either the primal dissipation potential *ψ*(***x, J***), with the state-change rate ***J*** as its variable, or the dual dissipation potential *ψ*^*^(***x***, *ξ*), with the thermodynamic force *ξ* as its variable.

For a standard passive dissipation structure, the instantaneous evolution rate at a fixed state ***x*** is determined by minimizing

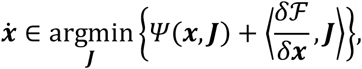

over admissible state-change rates ***J***. The two terms represent the dissipation cost and the rate of free-energy change, respectively. Under the standard convexity assumptions, this minimization principle is equivalent to the generalized gradient-flow relation in Eq. (1) through Legendre– Fenchel duality.

In this paper, we mainly use the dual dissipation potential because it directly yields the corresponding flux from the thermodynamic force, without explicitly solving the minimization problem at each instant. Unless otherwise stated, dissipation potential hereafter refers to the dual dissipation potential. The shifted coupling structure introduced in Section 5.2 is not a standard passive dissipation potential and is therefore interpreted separately as an active variational coupling.

### 3.3 Deriving dynamical equations from variational structures

Variational modeling based on the energy–dissipation principle proceeds in four steps:

1. Define the state variables: choose the probability distributions needed to describe the system (here, the gamete distribution ***g*** and the individual distribution ***ρ***);
2. Define the free energy ℱ: specify the equilibrium bias and driving forces of the system;
3. Choose the dissipation potential *ψ*^*^: characterize the kinetic response of the mechanism to the driving force (how the system moves);
4. Derive the dynamics: take the subdifferential with respect to the thermodynamic force to obtain the dynamical equation 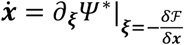

Section 4 applies this procedure to mutation, recombination, and selection separately, and Section 5 combines them into the complete model. For ease of reading, **Table 1** summarizes the main notation used throughout the paper.

**Table 1:** Summary of the main notation.

| notation | description |
| --- | --- |
| $\{0,1\}^L$ | haplotype space ($L$ biallelic loci) |
| $h$ | specific haplotype, $h \in \{0,1\}^L$ |
| $\Delta_{2^L-1}$ | probability simplex |
| $g_h$ | gamete state: frequency of haplotype $h$ in the gamete pool |
| $\rho_h$ | individual state: frequency of haplotype $h$ in the population of individuals |
| $\pi_h$ | mutation-equilibrium reference distribution |
| $\mathbf{g}, \mathbf{\rho}, \mathbf{\pi}$ | haplotype-frequency vectors |
| $\mathcal{F}$ | genetic free energy |
| $U(h)$ | selection potential $U$, interpreted as a fitness landscape |
| $\xi_{\mathbf{g},h}$ | thermodynamic force in the gamete space |
| $\xi_{\mathbf{\rho},h}$ | thermodynamic force in the individual space |
| $\Psi_{\text{mut}}^*$ | dual dissipation potential for mutation |
| $\Psi_{\text{rec}}^*$ | dual dissipation potential for recombination |
| $\Psi_{\text{sel}}^*$ | Fisher–Rao dual dissipation potential for selection |
| $\Psi_c^*$ | shifted coupling dissipation potential |
| $J_{\text{mut}}$ | mutation flux |
| $J_{\text{rec}}$ | recombination flux |
| $J_{\text{sel}}$ | selection flux |
| $J_c$ | coupling flux |
| $J_{\mathbf{g}}, J_{\mathbf{\rho}}$ | internal flux in the gamete / individual space |
| $\gamma$ | coupling strength |
| $N_e$ | effective number of haploid gene copies (twice the diploid effective population size when applicable) |

## 4 Variational Modeling of Mutation, Recombination, and Selection

Following the steps of Section 3, we build variational models for the gamete subsystem and the individual subsystem, respectively. Each subsystem consists of three components: a free energy, a dissipation potential, and the resulting dynamical equation. Detailed proofs are given in Appendix B.

### 4.1 Variational model of mutation and recombination at the gamete stage

The gamete subsystem is described by the gamete distribution ***g*** and incorporates two evolutionary processes: mutation and recombination. In the absence of selection, we show that these processes share a common free energy while being governed by distinct dissipation potentials, yielding a unified variational representation.

#### 4.1.1 Free energy of the gamete subsystem

We define the free energy of the gamete subsystem as

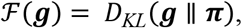

a Kullback–Leibler (KL) divergence that measures the deviation of the gamete distribution ***g*** from its reference equilibrium distribution ***π***, or equivalently, the relative entropy of ***g*** with respect to ***π***. Its variational derivative provides the thermodynamic forces for mutation and recombination. Here, the reference equilibrium distribution ***π*** is determined by the mutation rates and the detailed-balance condition.

##### Reference equilibrium distribution

On the full haplotype space {0,1}^*L*^, assuming that mutation acts independently across loci and satisfies detailed balance, the reference equilibrium distribution factorizes into a product of per-locus equilibrium distributions. For a biallelic locus, let *π*^(*l*)^ denote the equilibrium distribution at locus *l*, 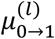 the forward mutation rate and 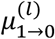 the backward mutation rate; the detailed-balance condition gives

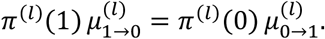

Thus, the reference equilibrium distribution is determined by the mutation rates; when the forward and backward mutation rates are equal, 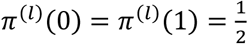.

The equilibrium frequency for the haplotype *h* is

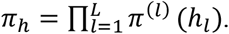

Here, *π*^(*l*)^ denotes the allele-frequency distribution at locus *l* under mutation equilibrium, with *π*^(*l*)^(0) and *π*^(*l*)^(1) denoting the equilibrium frequencies of the two alleles. The corresponding marginal distribution of ***g*** at locus *l* is denoted by *g*^(*l*)^, with 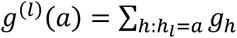.

##### Information decomposition and free-energy structure

Under the assumptions of independent mutation across loci and detailed balance, a key property of the relative entropy *D*_*KL*_(***g*** ∥ ***π***) is that it can be decomposed into marginal and association components:

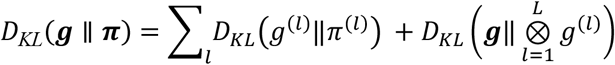

The first term measures deviations of the marginal allele frequencies from their mutation-equilibrium values. The second term, denoted by ***I***(***g***), measures multilocus statistical dependence, also known as total correlation. For two loci, ***I***(***g***) is an information-theoretic measure of linkage disequilibrium (LD) and vanishes exactly at linkage equilibrium. ***I***(***g***) also corresponds to the negative of the normalized entropy defined in Akin’s geometrical study of population genetics [31]. Relative-entropy dissipation toward the product of one-locus marginals has also been established for nonlinear recombination models [32]. The derivation of the decomposition is given in Appendix A.

##### Thermodynamic forces

The thermodynamic force in the gamete space is given by the negative variational derivative of the free energy with respect to ***g***:

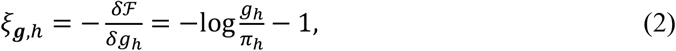

The vector *ξ*_***g***_ provides the thermodynamic driving force for mutation and recombination in the gamete space. In the absence of mutation, the reference distribution ***π*** is no longer defined by mutation equilibrium. For pure recombination, the relevant free-energy component is instead the total correlation ***I***(***g***) = *D*_KL_(***g*** ∥ ⊗_*l*_ *g*^(*l*)^), and the corresponding thermodynamic force is

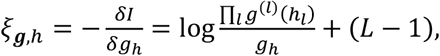

where the constant term (*L* − 1) cancels in the force differences entering the recombination fluxes and therefore does not affect the dynamics.

#### 4.1.2 Dissipation potentials for mutation and recombination

Both mutation and recombination can be represented as reversible reaction networks at the level of haplotype frequencies, and their dissipation potentials are constructed using the gradient-flow theory for reaction networks [29]. In this formulation, mutation corresponds to reversible jump reactions *h*^′^ ⇌ *h* between haplotypes, with first-order mass-action kinetics; recombination corresponds to binary reactions between pairs of haplotypes, (*h*_*A*_, *h*_*B*_) ⇌ (*h*_*C*_, *h*_*D*_), with second-order mass-action kinetics. Maas and Mielke [29] developed generalized gradient structures for detailed-balanced reaction networks based on reactant and product activities. Following their approach, we construct quadratic dual dissipation potentials with logarithmic-mean mobilities, whose gradient flows recover the corresponding classical mutation and recombination fluxes.

For mutation, detailed balance determines the reference distribution *π*. For recombination, the same product reference distribution satisfies detailed balance for each exchange event because recombination preserves the allele count at every locus; consequently, its contribution cancels from the event force.

Mutation and recombination share the same gamete free energy and thermodynamic force, but have distinct kinetic structures and therefore require different dissipation potentials.

##### Mutation dissipation potential

Mutation is a linear jump process between haplotypes. For a mutation-rate matrix *μ*_*ij*_ (from haplotype *i* to *j*), the classical mutation equation is

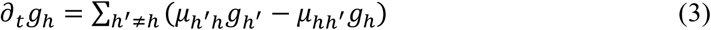

This is the standard representation of the mutation generator in reaction networks [28,29]. Under the detailed-balance condition, we define the equilibrium flux 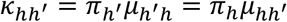, which guarantees a symmetric mobility. This logarithmic-mean construction is consistent with the entropy-gradient-flow representation of reversible finite Markov chains [33]. The associated mutation dissipation potential follows the standard construction based on a logarithmic-mean mobility:

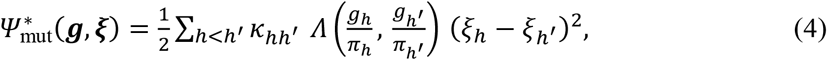

where 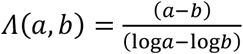is the logarithmic mean, with *Λ*(*a, a*) = *a* by continuity, and the sum is over unordered haplotype pairs *h* < *h*^′^ (each undirected edge counted once). The logarithmic-mean identity provides the key step connecting the variational representation to the standard mutation flux: using the gradient-flow relation 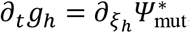, taking the subdifferential of the dissipation potential with respect to the thermodynamic force gives the classical mutation generator (Eq. 3). The detailed derivation is given in Appendix B.1.

##### Recombination dissipation potential

Recombination reshuffles genetic material between parental haplotypes during meiosis, changing multilocus association patterns while preserving marginal allele frequencies. A recombination event is written as (*h*_*A*_, *h*_*B*_) ↔ (*h*_*C*_, *h*_*D*_): the parental and recombinant haplotype pairs contain the same alleles but differ in their linkage phase. This defines a reversible binary reaction with second-order mass-action kinetics:

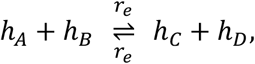

where *r*_*e*_ is the recombination rate per event. The corresponding reactant and product activities are

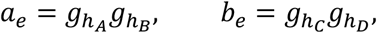

and the thermodynamic driving force is

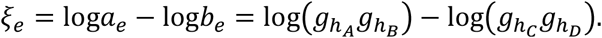

For arbitrary haplotype force variables, the event force is *ξ*_*e*_ = ∑_*h*_ *v*_*he*_ *ξ*_***g***,*h*_, with stoichiometric coefficients defined below. The recombination dissipation potential is quadratic in these event forces, with logarithmic-mean mobility *Λ*(*a*_*e*_, *b*_*e*_). The sum in Eq. (5) runs over distinct unordered exchange channels *e*, and *r*_*e*_ includes the required combinatorial factor. At the thermodynamic forces, the contribution of the product reference distribution ***π*** cancels because each event preserves allele counts, so the event force reduces to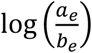 (Appendix B.2).

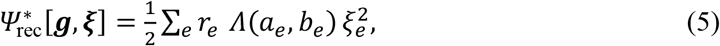

From the gradient flow 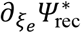, one obtains the per-event flux

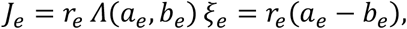

where the last equality follows from the logarithmic-mean identity. The stoichiometric coefficient *v*_*he*_ describes the net change in haplotype *h* caused by event *e*: it is +1 for a produced haplotype, −1 for a consumed haplotype, and 0 otherwise. Summing over all events gives the classical event-flux form:

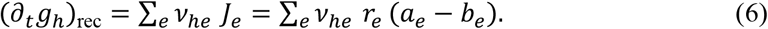

This is the standard stoichiometric flux representation of the recombination reaction network. In the pure-recombination limit without mutation and selection, for a genome comprising *L* loci, the dynamics reduce to the classical multilocus continuous-time recombination equation

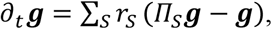

*S* indexes a chromosome partition, and *Π*_*S*_ denotes the corresponding recombination projection. This formulation is equivalent to the multilocus pure-recombination ordinary differential equations (ODEs) established in [21,31] (derivation in Appendix B.2).

#### 4.1.3 Dynamical equation of the gamete subsystem

Substituting the gamete free energy and the two dissipation potentials into the generalized gradient flow form (Eq. 1, Section 3.1) yields the dynamical equation of the gamete subsystem. With the thermodynamic 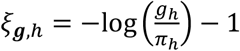 (Eq. 2) and the logarithmic-mean identity (Appendix B), the mutation and recombination fluxes reduce to their classical forms, and the gamete subsystem dynamics read

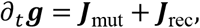

where 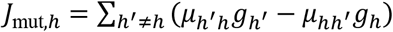 is the mutation flux (Eq. 3) describing local probability transfer between haplotypes, and *J*_rec,*h*_ = ∑_*e*_ *v*_*he*_ *r*_*e*_(*a*_*e*_ − *b*_*e*_) is the recombination flux (Eq. 6), obtained by summing the pairwise exchange-event fluxes. The coupling to the individual subsystem is introduced in Section 5. The derivations of these fluxes are given in Appendices B.1 and B.2.

The resulting formulation combines mutation and recombination through distinct dissipation potentials acting on the same gamete free energy, while preserving their classical kinetic forms.

### 4.2 Variational model of selection dynamics at the individual stage

The individual subsystem takes the individual-level haplotype distribution ***ρ*** as its state variable. The deterministic selection dynamics have a well-established gradient-flow structure under the Shahshahani metric [21,28]. Here, we incorporate this established structure into our variational framework, while genetic drift is treated separately as a stochastic extension in Section 6.

#### 4.2.1 Selection free energy

Selection is represented by a state-independent haplotype fitness potential *U*: {0,1}^*L*^ → ℝ, which assigns a selective advantage or disadvantage to each haplotype. Throughout the present formulation, *U*_*h*_ denotes the haplotype fitness potential. The free energy of the individual subsystem is therefore defined as the negative expected fitness,

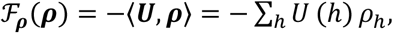

Thus, selection specifies the evolutionary driving force, whereas the dissipation potential determines the kinetic response to this force. This selection term should be distinguished from the mutation-related relative-entropy term: it represents fitness-driven selection rather than an equilibrium free energy associated with mutation. The thermodynamic force in the individual space is

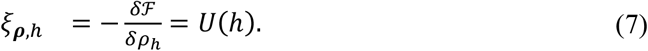

In the additive-selection models considered in our numerical experiments, *U*(*h*) = ∑_*l*_ *s*_*l*_ *b*_*h,l*_, where *s*_*l*_ is the per-locus selection coefficient, and *b*_*h,l*_ ∈ {0,1} indicates the allele carried by haplotype *h* at locus *l*.

#### 4.2.2 Selection dissipation potential: the Fisher–Rao geometry

Selection changes haplotype frequencies and can be represented geometrically on the probability simplex. We use the Shahshahani metric, equivalent to the discrete Fisher–Rao metric, to represent the selection dynamics on the simplex [21,34]. This geometric structure recovers the classical replicator equation and also determines the covariance matrix of the Langevin noise in the finite-population stochastic extension (Section 6).

The dual dissipation potential has a concise quadratic form on the discrete probability simplex 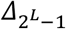:

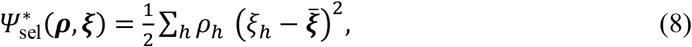

where 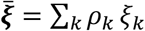 is the ρ-weighted mean thermodynamic force.

#### 4.2.3 Selection dynamical equation

From the generalized gradient-flow form (Eq. 1), the selection dynamics is obtained by taking the subdifferential of the selection dissipation potential (Eq. 8) with respect to *ξ*_***ρ***_ (derivation in Appendix B.3). Substituting the thermodynamic force *ξ*_***ρ***,*h*_ = *U*(*h*) (Eq. 7) gives the explicit selection dynamics (without yet considering the coupling to the gamete subsystem, which is introduced in Section 5):

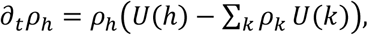

This is the classical replicator equation [35]: the frequency of each haplotype changes in proportion to its excess fitness. Subtracting the population-average force ensures conservation of total probability and yields this excess-fitness form. The resulting flow corresponds to the Shahshahani gradient flow on the discrete probability simplex.

## 5 The Gamete–Individual Two-State System

### 5.1 Coupling the two variational subsystems

Section 4 developed the gamete and individual subsystems separately, with each evolutionary process represented by its corresponding driving force and dissipation structure. To obtain a unified model, the two subsystems must be dynamically connected while preserving their distinct variational structures. We therefore introduce a shifted coupling potential that links the gamete and individual states without allowing the driving force of one stage to directly drive dissipation at the other stage.

### 5.2 Active coupling through a shifted dissipation potential

The coupling in our model provides a shifted dissipation structure that dynamically connects the two subsystems rather than representing an additional evolutionary process. Because the two states have different free-energy contributions and thermodynamic forces, we introduce a state-dependent active shift into a quadratic coupling potential centered by the mobility-weighted mean:

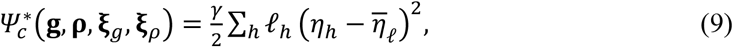

where *γ* > 0 is the coupling strength, *l*_*h*_ = *Λ*(*g*_*h*_, *ρ*_*h*_) is the logarithmic-mean mobility, *η*_*h*_ = *ξ*_*g,h*_ − *ξ*_*ρ,h*_ + *A*_*h*_ is the shifted coupling force, and 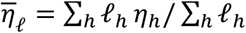 is its mobility-weighted mean. The state-dependent active shift *A*_*h*_(***g, ρ***) is defined as

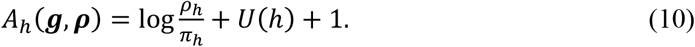

The shift reconciles the thermodynamic forces of the two subsystems. Subtracting the weighted mean ensures that the potential is invariant under independent constant shifts of these forces and that its coupling flux conserves probability. At fixed states and active shift, differentiation gives 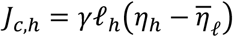. At the thermodynamic forces of Eqs. (2) and (7), the shifted force becomes

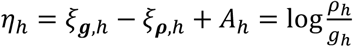

By the logarithmic-mean identity 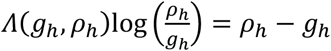 and probability normalization,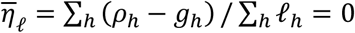. Differentiating the coupling potential with respect to the thermodynamic forces yields the coupling flux (see Appendix B.4):

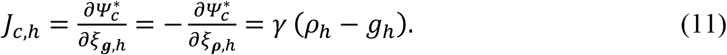

Thus, the coupling flux is directly proportional to the state difference (*ρ*_*h*_ − *g*_*h*_), providing relaxation toward consistency without introducing a coupling term in the free energy. The shifted potential remains an active variational representation, rather than a standard passive dissipation potential.

### 5.3 Complete two-state variational dynamics

The resulting model is a coupled variational system that combines passive dissipation structures with an active coupling shift.

Adding the dual dissipation potentials of the two subsystems and the shifted coupling potential gives the combined dual potential:

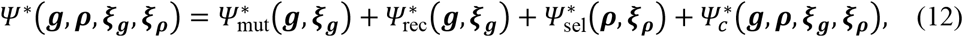

Adding the free energies of the two subsystems gives the total genetic free energy:

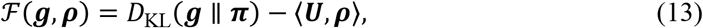

where ℱ _***g***_(***g***) = *D*_KL_(***g*** ∥ ***π***) is the internal free energy of the gamete pool (Section 4.1.1) and ℱ _***ρ***_(***ρ***) = −⟨***U, ρ***⟩ is the selection free energy in the individual space (Section 4.2.1). Each defines its corresponding thermodynamic force; gamete–individual consistency is enforced by the coupling dissipation potential 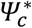 (Eq. 9).

The three passive dissipation potentials represent mutation, recombination, and selection, while the shifted coupling potential dynamically connects the two states. Mobility-weighted centering ensures that the coupling flux preserves probability normalization for arbitrary force variables without modifying the subsystem free energies or their thermodynamic forces. The complete dynamics follow from the subdifferential of the combined dual potential.

Complete two-state variational dynamics. Combining the three passive dissipation potentials with the shifted coupling structure gives the following coupled variational system, in which ∂_*ξ*_*ψ*^*^ denotes the subdifferential of *ψ*^*^ with respect to the force variable *ξ*:

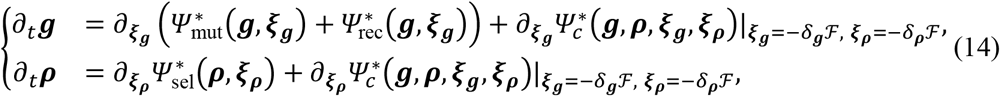

Equivalently, the dynamics can be written in terms of the internal and coupling fluxes:

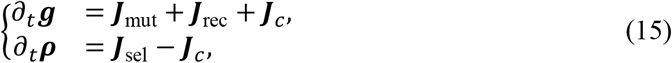

***J***_***g***_ = ***J***_mut_ + ***J***_rec_ is the internal flux in the gamete space (mutation and recombination; Eqs. 3 and 6), and ***J***_***ρ***_ = ***J***_sel_ is the internal flux in the individual space (selection); the internal fluxes are given by the subdifferentials of the corresponding modules, and the coupling flux is ***J***_*c*_ = *γ*(***ρ*** − ***g***) (Eq. 11). Note that ***J***_*c*_ is an active internal consistency flux between the two states, rather than an additional evolutionary process; it enters the two state equations with opposite signs, driving ***g*** and ***ρ*** toward consistency. Biological processes not explicitly modeled here, such as gamete-level selection or post-fertilization mutation in early embryos, could be introduced as additional subsystem-specific fluxes in future extensions. With mobility-weighted centering, the coupling flux automatically satisfies ∑_*h*_ *J*_*c,h*_ = 0 for arbitrary force variables, so both distributions remain normalized without a separate tangential projection.

Substituting the module-specific subdifferentials gives the complete expanded form. Gamete dynamics (mutation + recombination + coupling flux):

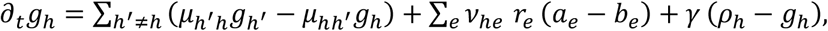

where the first term is the classical mutation generator (Eq. 3), the second is the classical pair-exchange flux, and the third is the coupling flux pulling ***g*** toward ***ρ***.

Individual dynamics (selection + coupling flux):

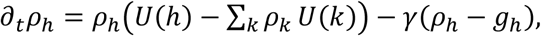

where the first term is the selection-driven Shahshahani flow (the subdifferential of the Fisher–Rao dissipation potential, with *ξ*_***ρ***,*h*_ = *U*(*h*)), and the second term is the coupling exchange flux pulling ***ρ*** toward ***g***.

The complete two-state system has the following key structural properties:

1. Probability conservation: the subdifferentials of 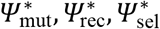 automatically lie in the tangent space of the simplex; mobility-weighted centering gives 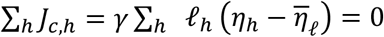 for arbitrary force variables. At the thermodynamic forces, the coupling flux ***J***_*c*_ = *γ*(***ρ*** − ***g***) satisfies ∑_*h*_ *J*_*c,h*_ = *γ* ∑_*h*_ (*ρ*_*h*_ − *g*_*h*_) = 0, so probability conservation holds automatically without a separate tangential projection.

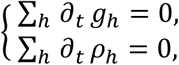
2. Modular structure preservation: the internal dynamics of each subspace (***J***_***g***_, ***J***_***ρ***_) preserve their own variational structures. Coupling is carried only by ***J***_*c*_, and this exchange flux does not change the total probability ∑_*h*_ (*g*_*h*_ + *ρ*_*h*_).

The total free energy ℱ is not generally a Lyapunov functional of the coupled system: the three passive modules dissipate it, whereas the active coupling contribution may have either sign. In single-process limits, the corresponding free energy recovers the usual monotone dissipation structure. This point was also discussed in Section 3.1.

### 5.4 Strong-coupling limit and connection to classical models

The coupling strength *γ* controls the stiffness of the consistency constraint between the two state variables. When *γ* → ∞, the coupling rapidly drives the state difference ***ρ*** − ***g*** toward zero, so that the dynamics approach the consensus manifold ***g*** = ***ρ*** = ***f***, and the two-state system reduces to a single-state system with reduced free energy ℱ _eff_(***f***) = *D*_KL_(***f*** ∥ ***π***) − ⟨***U, f***⟩, where ***f*** is the consensus state variable. The restriction of the total free-energy functional to the consensus manifold is this reduced free energy; however, Eq. (16) is obtained by singular reduction of the two-state dynamics, not by applying all dissipation potentials to the gradient of this restricted functional.

Under the consensus-state constraint, 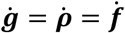. When the two equations in the dual-state flux form (Eq. 15) are added, the coupling flux ***J***_*c*_ cancels exactly, yielding the reduced single-state dynamical equation:

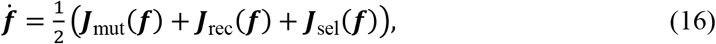

where ***J***_mut_(***f***), ***J***_rec_(***f***), and ***J***_sel_(***f***) are the mutation, recombination, and selection fluxes reduced to the consensus state, respectively. The consensus constraint reduces the state space, while the reduced dynamical equation is obtained by combining the two subsystem fluxes; this produces the factor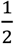 in the reduced deterministic dynamics. The factor 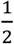 originates from the two stages evolving simultaneously: maintaining the constraint 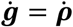requires the coupling flux to adjust to 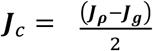, i.e., 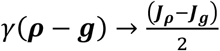 remains bounded and carries the constraint force. After the time rescaling *τ* = *t*/2, the factor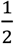 in Eq. (16) can be removed without changing the steady state. The detailed derivation is given in Appendix B.5.

The single-state dynamical equation (Eq. 16) directly relates the strong-coupling dynamics to Akin’s vector-field formulation[31], while the individual process-specific vector fields here are derived from their corresponding dual dissipation potentials.

For a single biallelic locus (*L* = 1), the consensus state ***f*** can be represented by a single allele frequency, denoted by *p*; the recombination flux vanishes, and the mutation and selection fluxes take their standard forms *J*_mut_ = *u*(1 − *p*) − *vp* and *J*_sel_ = *s p*(1 − *p*), so the single-state equation reads explicitly

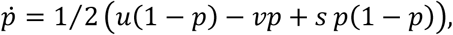

which is the classical mutation–selection equation (after the rescaling *τ* = *t*/2).

### 5.5 Considerations for extending the framework to diploid populations

This paper takes haploid populations as the default setting. For randomly mating diploid populations, the reproduction map generalizes to gamete fusion *T*(***g***) = ***g*** ⊗ ***g***, with random union of gametes generating zygotes in Hardy–Weinberg proportions. The haplotype distribution tracked in the current haploid formulation corresponds, in the diploid setting, to the marginal haplotype distribution induced by the diploid genotype distribution. When fitness depends on diploid genotype, the state must generally be extended from haplotype frequencies to genotype frequencies; the gamete–individual reproduction map then becomes an explicit gamete-fusion map, and the Fisher–Rao structure must be formulated on the resulting genotype simplex. For a single biallelic locus, for example, genotype-specific fitness can be specified by *W*_*AA*_, *W*_*Aa*_, and *W*_*aa*_, with the corresponding selection free energy expressed in terms of genotype frequencies. Selection involving dominance or epistasis, or non-random mating, can cause genotype frequencies to depart from Hardy–Weinberg proportions, so they cannot generally be recovered uniquely from haplotype marginals, requiring an expanded genotype-level state representation.

## 6 Stochastic Extension for Genetic Drift

Unlike mutation, recombination, and selection, genetic drift is not represented by a deterministic dissipation potential in our model. Instead, finite-population sampling fluctuations are incorporated as a stochastic extension of the deterministic coupled variational system, yielding a finite-population stochastic differential equation.

Motivated by finite-population sampling at the two life-cycle transitions (**Fig. 1**), we introduce Wright–Fisher-type noise in both state variables as a phenomenological stochastic extension.

### 6.1 Finite-population stochastic extension

Let the deterministic dynamics be given by the coupled variational system of Section 5.3. To account for finite-population sampling fluctuations, we add Langevin noise terms to both state equations, interpreted in the Itô sense:

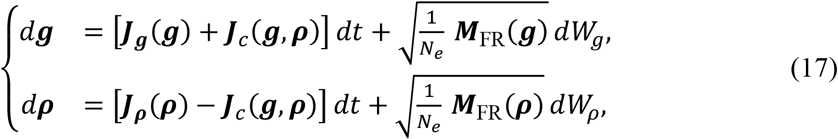

where ***J***_***g***_ = ***J***_mut_ + ***J***_rec_ and ***J***_***ρ***_ = ***J***_sel_ are the internal fluxes of the coupled system (Section 5.3), and *W*_*g*_ and *W*_*ρ*_ are independent standard Wiener processes. *N*_*e*_ is the effective number of haploid gene copies. The two noise terms represent finite-population sampling fluctuations associated with the two state transitions, rather than additional deterministic evolutionary forces. The state-dependent noise covariance is proportional to the Fisher–Rao mobility matrix:

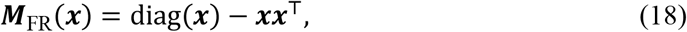

For a state ***x*** on the probability simplex, ***M***_FR_(***x***) has the standard multinomial sampling covariance structure of the Wright – Fisher process. For a single sampled gene copy, haplotype *h* has probability *x*_*h*_, giving diagonal entries *x*_*h*_(1 − *x*_*h*_) and off-diagonal entries −*x*_*h*_*x*_*k*_.

For the gamete state, ***M***_FR_(***g***) specifies the Wright– Fisher-type covariance structure; for the individual state, ***M***_FR_(***ρ***) specifies the corresponding covariance structure. The two sampling noises are assumed independent in the present formulation. This independence is a modeling assumption rather than a mathematical necessity; correlated sampling fluctuations could be incorporated through a joint noise covariance in future extensions. These state-wise noise terms are not derived here as the diffusion limit of two sequential multinomial sampling steps within a single generation. Using the same effective population size for both states is a simplifying assumption; the two stages may have different sampling intensities.

Thus, the stochastic extension adds finite-population sampling fluctuations without modifying the deterministic variational structure developed in Sections 3–5.

### 6.2 Connection to Wright–Fisher diffusion

In the strong-coupling limit, the two state variables converge to a common consensus state, and the stochastic two-state system reduces to a single stochastic process for the haplotype frequencies. With the consensus variable 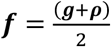 and the state difference ***y*** = ***ρ*** − ***g***, the coupling term drives the latter rapidly toward zero as *γ* → ∞. The two independent sampling contributions combine on the consensus manifold, yielding the reduced stochastic dynamics in the original time scale. Under the assumed independent, equal-strength Wright–Fisher-type sampling noises and the same time rescaling as in the deterministic reduction, the strong-coupling reduction recovers the standard Wright–Fisher diffusion covariance on the haplotype-frequency simplex. In the single-locus biallelic case, the corresponding Fokker–Planck equation is the classical Kimura diffusion equation. The pure-drift case follows by setting the deterministic evolutionary forces to zero. The detailed derivations are given in Appendix C.

## 7 Numerical Experiments

The theoretical framework is evaluated through three progressively complex experiments: Experiment 1 (Section 7.1) tests whether the variational dynamics recover the classical single-locus mutation–selection equation; Experiment 2 (Section 7.2) extends the analysis to a two-locus haplotype space and evaluates the evolution of multilocus probability distributions; and Experiment 3 (Section 7.3) integrates mutation, recombination, and selection in a three-locus system and evaluates the finite-population stochastic extension. Together, these experiments test whether the variational population genetics (VPG) framework developed above recovers classical population-genetic dynamics and extends to multilocus and finite-population stochastic settings. Deterministic trajectories are obtained by numerical integration of the derived dynamical equations, stochastic trajectories by Langevin integration, and finite-population benchmarks by SLiM forward simulations [13]. The mathematical notation numbers loci from 1; simulation labels use zero-based indexing (e.g., site 0, site 1, etc.).

### 7.1 Single-locus mutation–selection balance

We first test the VPG framework in the simplest genetic system—a single biallelic locus—where mutation and selection jointly determine the equilibrium allele frequency.

We consider 11 parameter sets: the nine combinations of the three symmetric mutation rates *u* = *v* = *μ* ∈ {3 × 10^−4^, 10^−3^, 3 × 10^−3^} and three selection coefficients *s* ∈ {0.005,0.01,0.05}, each initialized at the low frequency used for the main trajectory plot (**Fig. 2A**), together with two additional parameter sets initialized at *p*(0) = 0.99 (the two larger mutation rates paired with the middle selection coefficient), to test bidirectional convergence (**Fig. 2B**). The main trajectory plot (**Fig. 2A**) uses *μ* = 10^−3^ and *p*(0) = 0.01. The effective number of haploid gene copies is *N*_*e*_ = 2000 (1000 diploid individuals), and we perform 100 independent replicate simulations in SLiM under a Wright–Fisher model (detailed SLiM settings in Appendix D.1.2). The deterministic analyses compare the single-state variational population genetics (SVPG) and dual-state variational population genetics (DVPG) formulations, with coupling strength *γ* = 100 and the time rescaling *τ* = *t*/2 for the latter.

**Figure 2:**
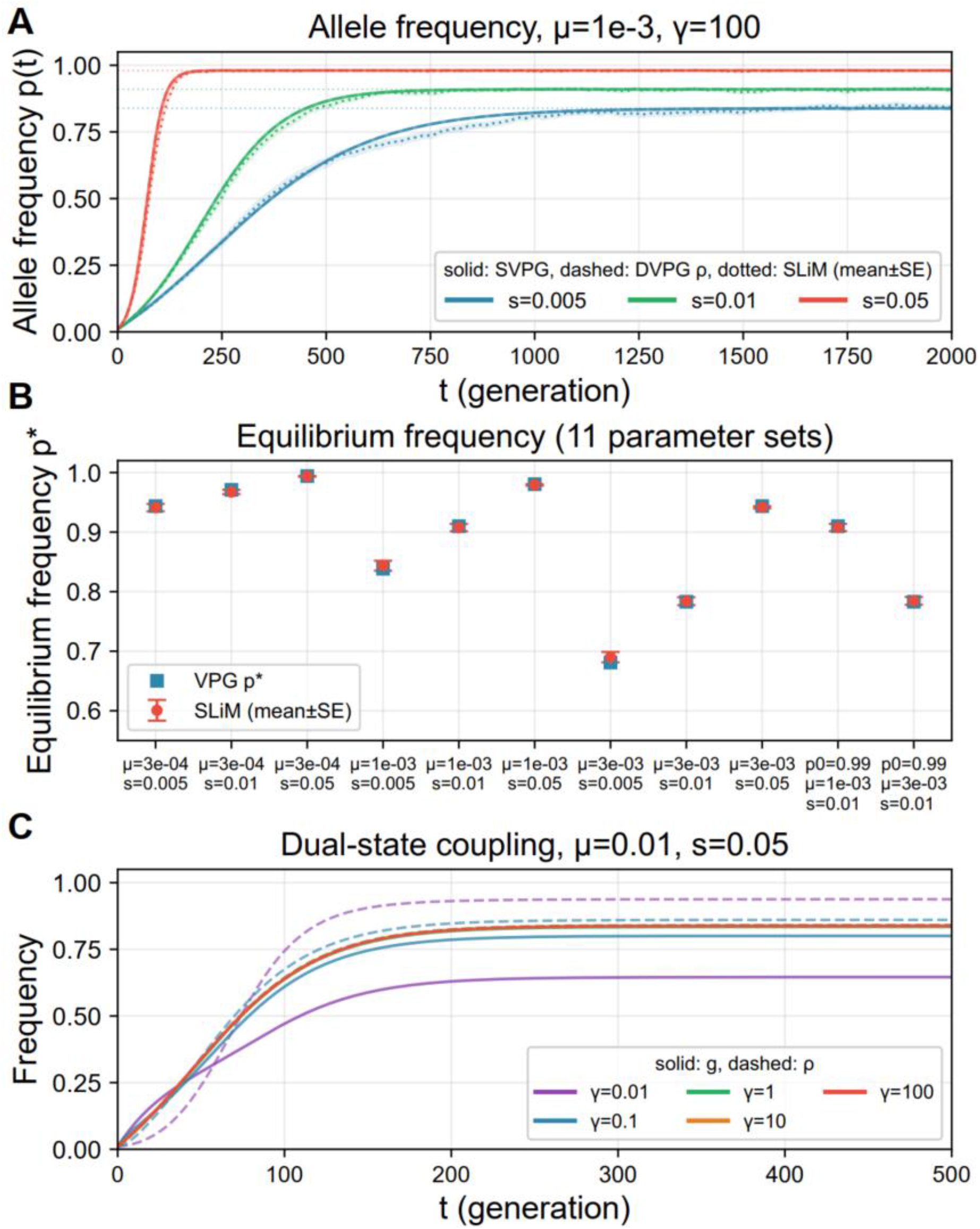
Numerical experiment for single-locus mutation–selection balance. **(A)** Allele frequency trajectories (μ = 10^−3^, p(t = 0) = 0.01, γ = 100): SVPG thin solid lines, DVPG **ρ** dashed lines (time rescaled as τ = t/2 to align with SVPG), SLiM mean dotted lines (shaded mean±SE). **(B)** Equilibrium frequency comparison: the VPG analytical equilibrium is consistent with the SLiM mean at t = 2000 for different parameter settings. **(C)** Coupling synchronization: allele frequencies AF(**g**) and AF(**ρ**) converge as γ increases (D_gρ_ = |g − ρ| ≈ 6.8 × 10^−5^ at γ = 100).

Across 11 parameter sets, SVPG trajectories and analytical equilibrium solutions are consistent with the SLiM simulations (**Fig. 2A,B**). DVPG converges to the same dynamics as SVPG under strong coupling, while increasing the coupling strength synchronizes ***g*** and ***ρ*** (**Fig. 2C**). These results confirm consistency between the variational formulation and classical single-locus mutation–selection dynamics.

### 7.2 Two-locus recombination–selection dynamics

We consider two biallelic loci (*L* = 2), with mutation turned off and selection acting only on the second locus (simulation site 1). The initial configuration has strong positive linkage disequilibrium: *p*_00_ = 0.45, *p*_11_ = 0.45, *p*_01_ = *p*_10_ = 0.05 (*D*(0) = 0.20). The parameter grid is *s* ∈ {0.005,0.05} × *r* ∈ {0.001,0.01}, with *γ* = 100, *N*_*e*_ = 2000 haploid gene copies (1000 diploid individuals), and 100 independent SLiM simulations under a Wright–Fisher model (detailed SLiM settings in Appendix D.2.2).

VPG closely tracks the four haplotype-frequency trajectories observed in SLiM, including the transient increase of recombinant haplotypes and the final enrichment of the favorable haplotype (**Fig. 3A**). LD decays with increasing recombination rate and is ultimately reduced toward linkage equilibrium, while the neutral locus—initially associated with the selected locus through positive LD—changes through genetic hitchhiking, and stronger recombination weakens this indirect response to selection (**Fig. 3B,C**). Thus, VPG jointly captures haplotype-frequency, allele-frequency, and LD dynamics.

**Figure 3:**
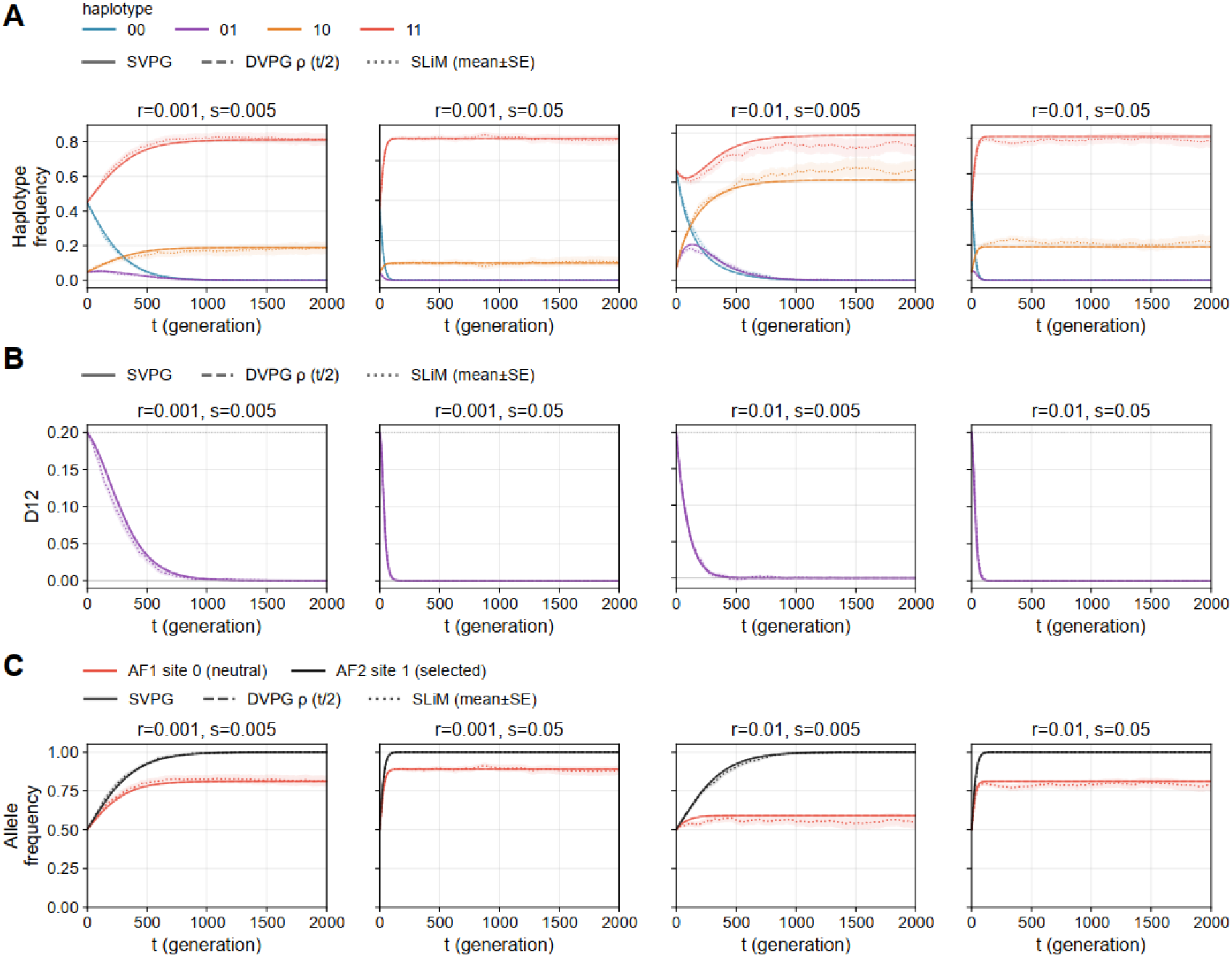
Numerical experiment for two-locus recombination–selection dynamics. Four parameter combinations: (r, s) = (0.001,0.005), (0.001,0.05), (0.01,0.005), (0.01,0.05). **(A)** Haplotype-frequency dynamics of the four haplotypes (SVPG thin solid, DVPG **ρ** thick dashed, SLiM dotted, mean±SE). **(B)** LD dynamics D_12_(t): LD decays faster at higher recombination rates. **(C)** Allele-frequency dynamics at the neutral and selected loci.

### 7.3 Three-locus integrated dynamics and stochastic extension

We consider three biallelic loci with symmetric mutation rates *u* = *v* = 0.001, locus-specific selection *s* = [+0.005,0, −0.005], and recombination rates *r*_12_ = 0.01, *r*_23_ = 0.05 on the two intervals. The initial configuration has high LD: *p*_000_ = *p*_111_ = 0.4. We perform 100 independent simulations in SLiM under a Wright–Fisher model for each of three diploid population sizes (500, 1000 and 3000 individuals; detailed SLiM settings in Appendix D.3.2). For the stochastic extension, an SVPG model with genetic drift (Langevin noise) is compared with SLiM in terms of the mean trajectories of allele frequencies and LD. For the stochastic numerical experiment, we use the single-state reduction as the finite-population benchmark, with the stochastic integration settings given in Appendix D.3.3.

VPG closely tracks the dynamics of all eight haplotypes, including enrichment of positively selected combinations, depletion of deleterious combinations, and recombination-driven restructuring of haplotype frequencies (**Fig. 4A**). The corresponding allele-frequency and LD trajectories are also consistent with the expected effects of selection and recombination (**Fig. 4B,C**). The stochastic extension gives replicate means and standard errors comparable to those from SLiM (**Fig. 4D,E,F**). These results provide a numerical consistency check for the Fisher–Rao stochastic extension in the tested multilocus setting.

**Figure 4:**
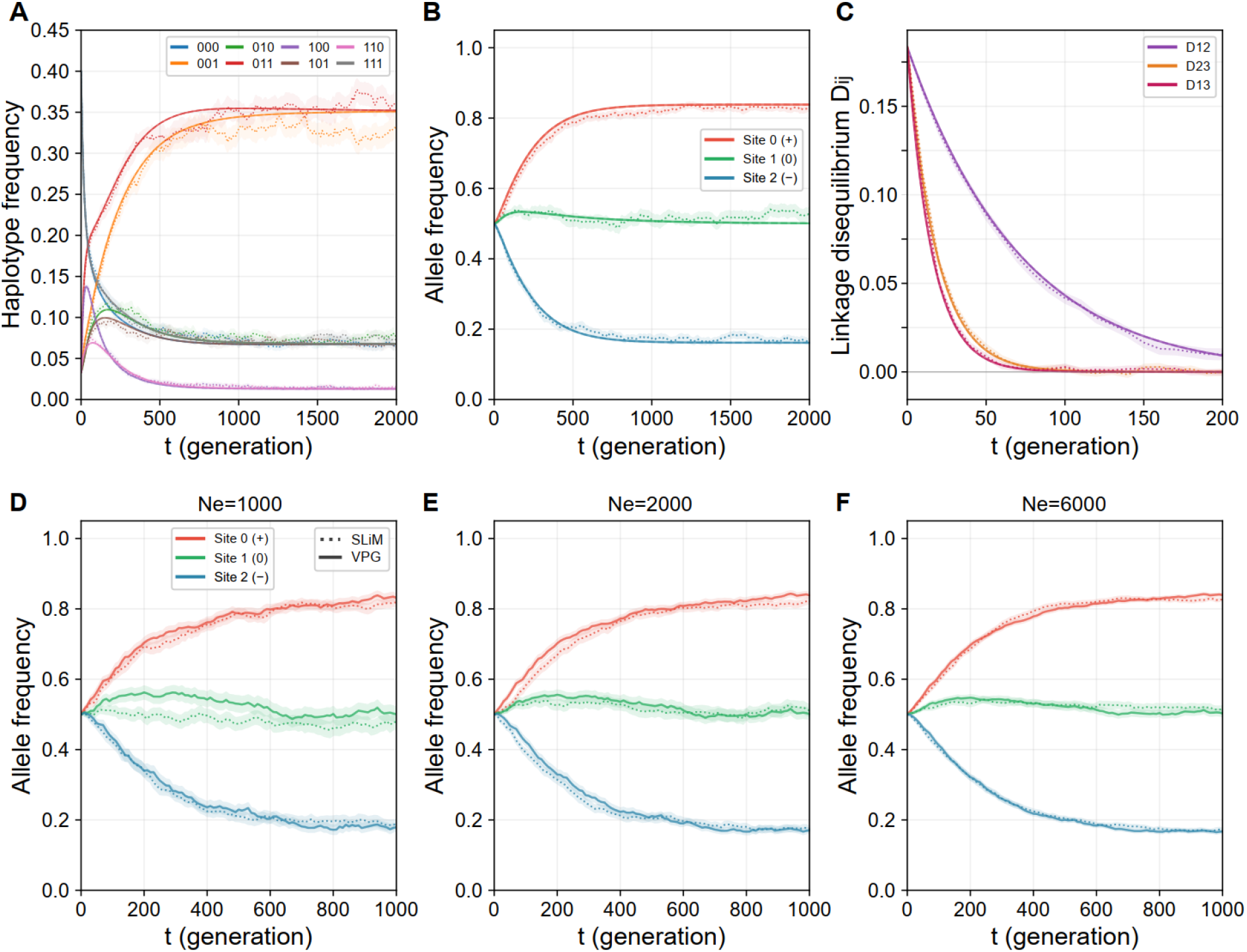
Numerical experiment for three-locus integrated dynamics and stochastic extension. **(A)** Dynamics of all 8 haplotype frequencies (SVPG thin solid lines, DVPG **ρ** thick dashed lines, SLiM dotted lines): haplotypes carrying the positively selected allele at locus 0 are enriched, haplotypes carrying the deleterious allele at locus 2 are strongly depleted, and recombination reshapes the haplotype structure. **(B)** Allele frequencies at the three loci: the frequency of the positively selected allele increases, the neutral allele remains approximately stable, and the negatively selected allele decreases. **(C)** Multilocus LD dynamics: D_12_ (r_12_ = 0.01) decays slowly, while D_23_ (r_23_ = 0.05) and D_13_ decay quickly—the LD decay patterns are consistent with the corresponding recombination rates. **(D-F)** Stochastic extension with three different N_e_ values (1000/2000/6000): comparison of AF between the SVPG model with genetic drift and SLiM (solid lines VPG, dotted lines SLiM, shaded mean±SE, both 100 replicates; first 1000 generations shown)—the stochastic extension gives comparable replicate means and standard errors.

### 7.4 Summary

Across the three experiments, VPG shows consistency with classical mutation–selection, multilocus haplotype/LD, and finite-population stochastic dynamics under the tested settings. Detailed simulation parameters and computational procedures are provided in Appendix D.

## 8 Discussion

### 8.1 Main contributions

This study develops a variational framework for modeling population genetic dynamics. The framework combines multiple evolutionary processes within a common mathematical formulation while preserving their distinct dynamical roles.

First, the framework provides a modular variational representation in which evolutionary processes are encoded by process-specific dissipation structures. In particular, mutation and recombination share a common gamete free energy but use distinct dissipation potentials that recover their classical fluxes. This allows multiple evolutionary processes to be combined within a common variational framework.

Second, we introduced a gamete–individual two-state representation that explicitly distinguishes successive life-cycle stages. Mutation and recombination act on the gamete state, whereas selection acts on the individual state, with coupling dynamically enforcing consistency between the two stage-specific distributions.

Third, we incorporated finite-population stochasticity phenomenologically through Wright–Fisher-type sampling noise in both state variables. Under the assumed independent, equal-strength noises and the same time rescaling as in the deterministic reduction, the strong-coupling limit recovers the standard Wright–Fisher diffusion covariance.

The framework was evaluated through three numerical experiments covering mutation–selection dynamics, recombination–selection dynamics, and their combined effects. These experiments demonstrate that the variational formulation remains consistent with the expected population genetic dynamics under the regimes considered.

### 8.2 Relation to classical population genetic models

The proposed framework is designed to remain consistent with established population genetic models rather than replace them. Under the assumptions considered here, the variational formulations recover the corresponding classical deterministic dynamics: the mutation, selection, and recombination components reduce to classical mutation-balance, replicator, and linkage-disequilibrium dynamics, respectively [21,28]. The stochastic formulation is connected to Wright– Fisher diffusion under the noise assumptions and time rescaling specified in Section 6.2 [36,37].

Akin [31] represented mutation, recombination, and selection as distinct vector fields on the haplotype-frequency space and defined the combined dynamics by directly adding these vector fields. Our model retains this process-wise modularity at the variational level, with each mechanism represented by a separate dual dissipation potential; in the strong-coupling limit, where the gamete and individual states coincide, the resulting single-state dynamics can be directly related to Akin’s combined vector-field formulation.

Combined evolutionary dynamics need not be gradient-like: selection–recombination models can exhibit cycles even with constant fitness [31], while selection–recombination–mutation models can sustain oscillations under certain non-independent mutation schemes [38]. Thus, the variational structures of individual processes do not by themselves imply a global gradient-flow structure for the coupled system.

### 8.3 Implications for differentiable population genetics

Automatic differentiation enables efficient computation of gradients through complex models, including high-dimensional systems of differential equations. Combined with deep learning, it has made gradient-based optimization increasingly practical for models with many parameters, and has been extended to scientific modeling through differentiable programming, such as differentiable physics, which combines differentiable programming with physics-based models [39].

A similar approach to population genetics could use gradients of predicted genetic variation with respect to evolutionary parameters for inference. The explicit mutation, recombination, and selection fluxes developed here provide a starting point for such differentiable population genetics, potentially enabling joint estimation of fine-scale mutation rates, selection coefficients, and recombination rates. Practical inference would also require methods to compare model predictions with observed genomic variation, distinguish the effects of different evolutionary parameters, and compute gradients reliably; a variational formulation alone does not ensure efficient inference. Implementing and evaluating these methods remains beyond the scope of this study.

### 8.4 Limitations and outlook

Several limitations remain. First, the current framework assumes panmixia, and the numerical tests cover only one to three loci with additive selection. A general, frequency-independent haplotype fitness potential can encode epistasis, but such interactions have not been evaluated here. Population structure, migration, diploid genotype-dependent fitness, and frequency-dependent selection require further development or validation.

Second, several modeling assumptions restrict the current scope of application. The present formulation relies on continuous-time dynamics and, for some variational constructions, detailed balance and independent-locus mutation assumptions. Relaxing these assumptions would allow the framework to accommodate more general scenarios of molecular evolution. Genetic drift is represented phenomenologically by independent, equal-strength Wright–Fisher-type noises at the gamete and individual stages. Applications to real population genomic data will require further refinement and validation of this stochastic extension, including assessment of demographic history and possible correlations between stages.

Finally, the full haplotype state space grows exponentially with the number of loci, so a direct full-state formulation cannot scale to large genomic regions. Future work should therefore develop hierarchical or reduced-order variational models that preserve the relevant population genetic dynamics and genetic associations while reducing the dimensionality of the state space, potentially drawing on model-reduction strategies from variational theories and statistical physics. These developments, together with empirical validation on real genomic datasets, will be important for extending the framework toward genome-scale applications.

## Appendix Appendix A: Decomposition of the Gamete Free Energy

This appendix derives the decomposition of the gamete free energy presented in Section 4.1.1. The decomposition relies on the factorization of the mutation-equilibrium reference distribution across loci.

We start from the KL divergence and insert the product of the marginal distributions ∏_*l*_ *g*^(*l*)^(*h*_*l*_) into both numerator and denominator:

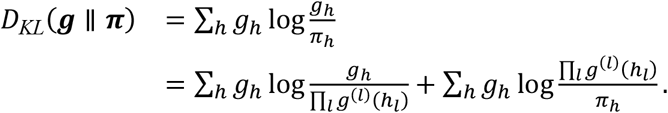

The first term is precisely the total correlation ***I***(***g***) = *D*_*KL*_(***g*** ∥ ⊗_*l*_ *g*^(*l*)^). For the second term, using the product structure of the reference distribution *π*_*h*_ = ∏_*l*_ *π*^(*l*)^(*h*_*l*_), we split the logarithm into a sum over loci:

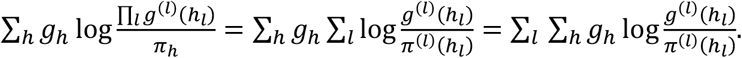

For fixed *l*, group the sum according to the allelic value *h*_*l*_: denote by 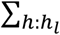 the sum over the remaining loci with *h*_*l*_ fixed; then

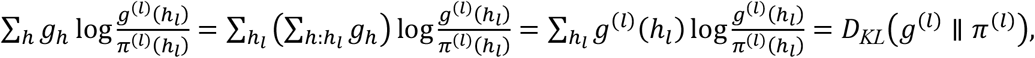

where the last step uses the definition of the marginal distribution 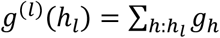. Combining the two terms gives

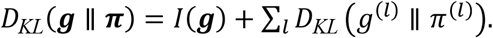

## Appendix B: Variational Derivations and Deterministic Reductions

This appendix gives the detailed variational derivations of the mutation, recombination, selection, and coupling fluxes introduced in Sections 4 and 5, together with the deterministic reductions to classical population genetic dynamics.

### B.1 Mutation flux derivation

We start from the mutation dissipation potential (Eq. 4), with equilibrium flux 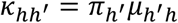 and logarithmic-mean mobility: 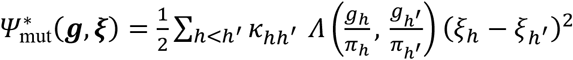. With recombination and coupling switched off and stochastic fluctuations absent, the gamete gradient flow reduces to the classical mutation generator.

Step 1. Free energy and thermodynamic force. Under pure mutation, the free energy is:

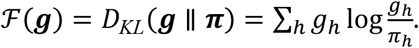

The variational derivative is:

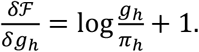

The thermodynamic force is:

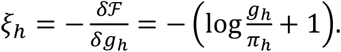

Step 2. Subdifferential. Taking the subdifferential with respect to *ξ*_*h*_:

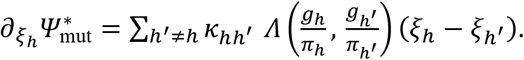

Step 3. Logarithmic-mean identity. Since 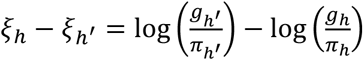, the classical logarithmic-mean identity gives:

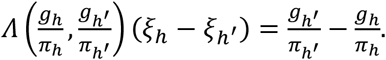

This identity holds for any ***π*** and does not require a uniformity assumption.

Step 4. Recovering the classical generator. Substituting the subdifferential and using detailed balance 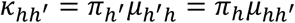:

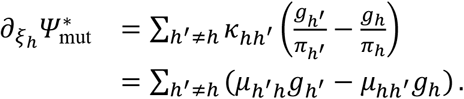

Hence 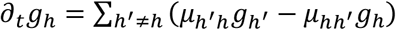, the classical mutation master equation.

### B.2 Recombination flux and multilocus reduction

We start from the recombination dissipation potential of the main text (Eq. 5). Its generalized gradient flow 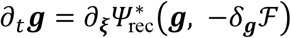 recovers the classical pair-exchange flux exactly. We then show how the event-level representation reduces to the classical multilocus recombination equation.

Step 1. Subdifferential of the dissipation potential. The recombination dissipation potential is constructed as (logarithmic mean + quadratic potential):

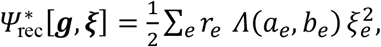

where 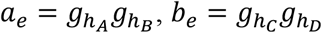, and the event force *ξ*_*e*_ is the stoichiometric combination of haplotype forces:

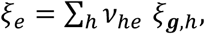

where *v*_*he*_ is the stoichiometric coefficient: +1 when event *e* produces haplotype *h*, −1 when it consumes *h*, and 0 when it does not involve *h*.

Taking the subdifferential with respect to *ξ*_***g***,*k*_ and applying the chain rule:

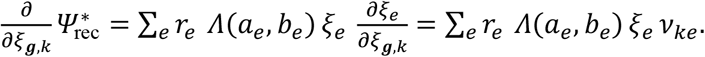

Step 2. Substituting the thermodynamic force. The thermodynamic force is 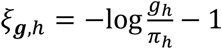.

The dynamics is formulated in terms of the gamete distribution ***g***; the thermodynamic force in the gamete space does not contain the selection potential *U* (*U* enters the free energy only through the individual distribution ***ρ***), so no contribution of *U* appears in *ξ*_*e*_. We substitute this expression into the event force and use ∑_*h*_ *v*_*he*_ = 0 (each event produces and consumes equal numbers of haplotypes) to eliminate the constant term. Because *π*_*h*_ = ∏_*l*_ *π*^(*l*)^(*h*_*l*_) and the reaction conserves the allele count at every locus, *logπ*_*h*_ is additive across loci and satisfies ∑_*h*_ *v*_*he*_*logπ*_*h*_ = 0. Thus, the contribution of ***π*** cancels exactly in *ξ*_*e*_:

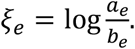

Step 3. Logarithmic-mean identity. Using the identity (which holds exactly with 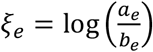:

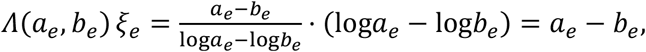

This holds exactly and does not require a weak-LD approximation.

Step 4. Flux recovery. Substituting this identity into the derivative gives

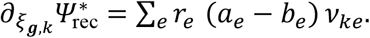

Thus, the flux on each event edge is *J*_*e*_ = *r*_*e*_ (*a*_*e*_ − *b*_*e*_) (with the sign determined by the convention for the stoichiometric coefficient *v*_*he*_), exactly recovering the classical pair-exchange flux.

Step 5. Multilocus reduction. In the pure-recombination limit, the event-level flux can be regrouped according to chromosome partition patterns. Let *S* ⊂ {1, …, *L*} denote the set of loci inherited from one parental haplotype, with *S*^*c*^ inherited from the other parent; we identify *S* and *S*^*c*^ as the same partition and sum over a canonical set of distinct bipartitions. Under this convention, each pair-exchange event is assigned to its associated canonical bipartition. For a fixed partition *S*, the recombination operator maps the haplotype distribution ***g*** to the product of its marginals on the two locus sets. Denoting this projection by *Π*_*S*_***g***, the net contribution of all events assigned to this partition *S* can be written as *r*_*S*_(*Π*_*S*_***g*** − ***g***), where *r*_*S*_ is the total rate associated with that partition. The event rates are defined to include the relevant multiplicities, so no additional combinatorial factor is introduced at the regrouping step. Summing over the canonical set of distinct bipartitions gives

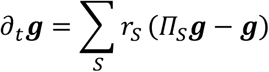

which is the standard continuous-time multilocus recombination equation. Thus, the event-level variational formulation is equivalent to the partition-based recombination dynamics under the above event and rate conventions.

### B.3 Fisher–Rao selection flux derivation

For the Fisher–Rao dissipation potential of the main text (Eq. 8), the partial derivative of the dual dissipation potential 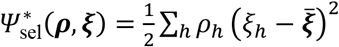with respect to 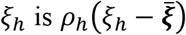, where 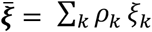

Computing the partial derivative:

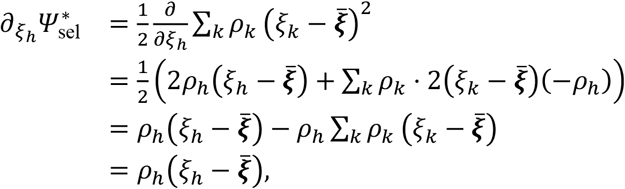

where we used 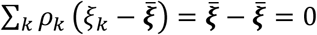.

### B.4 Coupling potential and flux derivation

We start from the shifted coupling potential of the main text (Eq. 9). For *g*_*h*_, *ρ*_*h*_, *π*_*h*_ > 0 and *γ* > 0, we hold the states, mobilities *l*_*h*_ = *Λ*(*g*_*h*_, *ρ*_*h*_), and active shift *A*_*h*_ fixed when differentiating with respect to the force variables. We write *S*_*l*_ = ∑_*h*_ *l*_*h*_ > 0, with *Λ*(*a, a*) = *a* by continuity.

Step 1. Nonnegativity and convexity. The coupling potential is nonnegative and convex in the force variables because it is a positive weighted sum of squares of affine functions. Its minimum value is zero and is attained when *η*_*h*_ is constant. However, the potential need not vanish when the unshifted forces are zero, so it represents active coupling rather than standard passive dissipation.

Step 2. Invariance under constant force shifts. Independent shifts **ξ**_*g*_ ↦ **ξ**_*g*_ + *c*_*g*_**1** and **ξ**_*ρ*_ ↦ **ξ**_*ρ*_ + *c*_*ρ*_**1** add the same constant *c*_*g*_ − *c*_*ρ*_ to *η*_*h*_ and 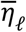. Their difference, and hence the potential and its force derivatives, are unchanged.

Step 3. Differentiation and probability conservation. Since 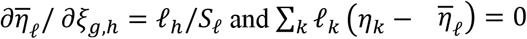, differentiation gives

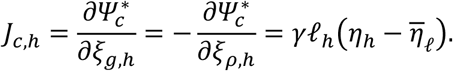

The coupling enters the two state equations with opposite signs and preserves the normalization of each distribution for arbitrary force variables:

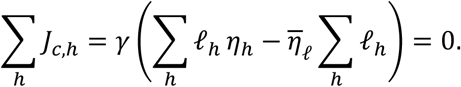

Step 4. Substitution of the thermodynamic forces. Substituting the thermodynamic forces (Eqs. 2 and 7) and the active shift (Eq. 10) gives

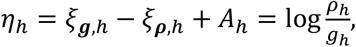

By the logarithmic-mean identity *l*_*h*_*η*_*h*_ = *ρ*_*h*_ − *g*_*h*_ and the normalization of both distributions, we obtain

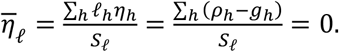

Consequently, the coupling flux is 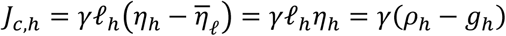, recovering Eq. (11).

The derivation above applies in the interior of the product simplex. Although logarithmic forces may be singular at the boundary, the expanded flux **J**_*c*_ = *γ*(**ρ** − **g**) extends continuously to the closed product simplex.

### B.5 Strong-coupling deterministic reduction

In the strong-coupling limit *γ* → ∞, the coupling term constrains the two states to the consensus manifold ***g*** = ***ρ*** = ***f***. Starting from the deterministic flux form,

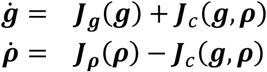

adding the two equations eliminates the coupling flux:

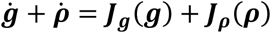

On the consensus manifold, ***g*** = ***ρ*** = ***f***, and hence

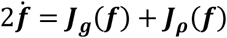

Therefore,

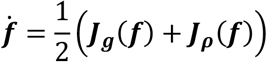

The factor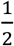 reflects the two parallel state equations. Introducing the rescaled time 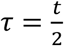 gives

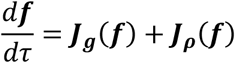

which gives the corresponding classical single-state population-genetic dynamics. The time rescaling changes only the parametrization of the trajectory and therefore does not change its equilibrium states.

## Appendix C: Stochastic Extension and Diffusion Reduction

This appendix gives the detailed derivation of the stochastic extension of Section 6 and of the diffusion reductions discussed in Section 6.2.

### C.1 Stochastic two-state dynamics

Let the state be ***X*** = (***g, ρ***), with total free energy

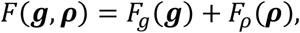

where *F*_*g*_(***g***) and *F*_*ρ*_(***ρ***) denote the gamete and selection free energies defined in Sections 4.1.1 and 4.2.1. The stochastic extension of the coupled variational system is obtained by adding finite-population sampling fluctuations to both state equations in the Itô sense:

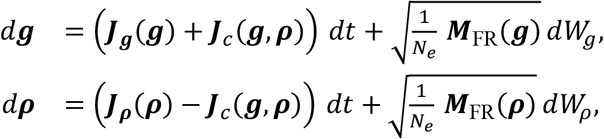

Here, *W*_*g*_ and *W*_*ρ*_ are independent standard Wiener processes, and the matrix square root is understood in the usual sense, so that the noise covariance in each state is

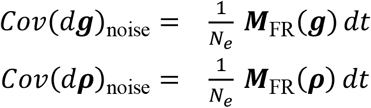

### C.2 Strong-coupling stochastic reduction

In the strong-coupling limit, the coupling term drives the two state variables toward a common consensus state. Introduce the consensus and difference variables

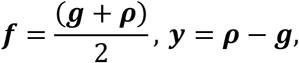

Step 1. Consensus variable. Adding the two stochastic state equations cancels the coupling flux exactly, giving

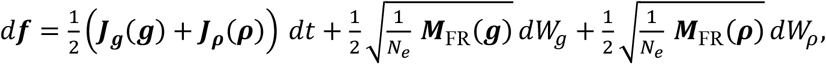

Step 2. Difference variable. Subtracting the two equations gives

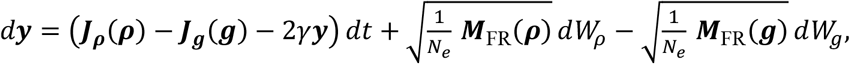

Thus ***y*** is a fast mean-reverting variable as *γ* → ∞, whereas ***f*** is the slow haplotype-frequency variable.

Step 3. Consensus limit and covariance. In the strong-coupling limit, ***y*** → 0, so that ***g, ρ*** → ***f*** and ***M***_FR_(***g***), ***M***_FR_(***ρ***) → ***M***_FR_(***f***). The stochastic increment of ***f*** therefore has covariance

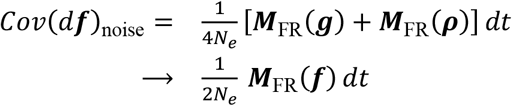

where the cross-covariance terms vanish because *W*_*g*_ and *W*_*ρ*_ are independent. Hence, on the consensus manifold,

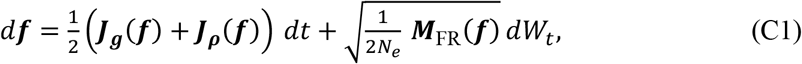

Step 4. Time rescaling. Under the time rescaling used in the deterministic reduction, 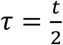, *dt* = 2*dτ*, and 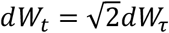, Eq. (C1) becomes

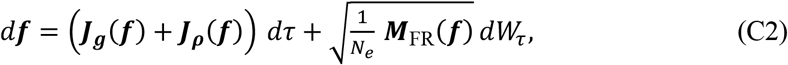

Thus, under the assumption of two independent sampling noises of equal strength and the same time rescaling, the strong-coupling stochastic reduction recovers both the classical deterministic rates and the standard Wright–Fisher sampling covariance.

### C.3 The pure-drift limit and the single-locus Kimura diffusion equation

In the pure-drift limit, with mutation, recombination, and selection switched off (***J***_***g***_ = ***J***_***ρ***_ = **0**), Eq. (C2) reduces to

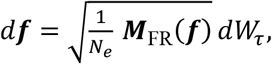

which, under the assumption of two independent sampling noises of equal strength and the same time rescaling, is the standard Wright–Fisher diffusion on the haplotype-frequency simplex. The corresponding Fokker–Planck equation for the probability density *q*(***f***, *τ*) is

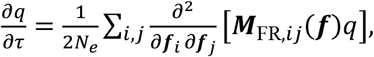

For a single biallelic locus, ***f*** can be represented by the allele frequency *p*, whose marginal diffusion coefficient is determined by *M*_FR_(*p*) = *p*(1 − *p*). Including the deterministic mutation– selection drift *b*(*p*) = *J*_mut_(*p*) + *J*_sel_(*p*), Eq. (C2) becomes

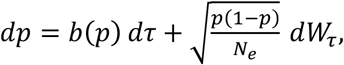

Its Fokker–Planck equation is

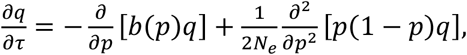

which is the classical Kimura diffusion equation.

## Appendix D: Numerical Methods and Supplementary Results

This appendix provides detailed simulation settings, numerical solution methods, and parameter-sweep results for the three numerical experiments in Section 7. For each parameter set, 100 independent SLiM forward simulations are performed under a Wright–Fisher model. Shaded regions indicate the mean ± SE across independent replicates.

### D.1 Supplementary material for Experiment 1: single-locus mutation– selection balance

#### D.1.1 Numerical solution

The deterministic ODEs are integrated with a time step of *dt* = 0.01 generations over *T* = 2000 generations. To account for the factor of one-half in the strong-coupling reduction, DVPG is integrated to *t* = 4000 in model time and plotted against the rescaled time *τ* = *t*/2, corresponding to 2000 generations on the SVPG time axis. SVPG is integrated using an explicit Euler scheme with per-step probability-simplex stabilization by clipping negative components followed by renormalization; this stabilization procedure is used only for numerical robustness and is not part of the continuous-time VPG dynamics. For DVPG, operator splitting with a matrix-exponential step is used for the linear coupling subsystem ***J***_*c*_ = *γ*(***ρ*** − ***g***), which preserves the mean state and causes the state difference to decay as *e*^−2*γdt*^. The mutation and selection fluxes are integrated using explicit Euler steps. Convergence is assessed by comparing the final frequency with the positive root of the corresponding quadratic equation 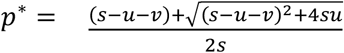. A deviation below 10^−5^ is considered converged. The main figure uses a coupling strength of *γ* = 100, for which *D*_*gρ*_ ≈ 6.8 × 10^−5^. The coupling-synchronization sweep in Fig. 2C uses *γ* ∈ {0.01,0.1,1,10,100}.

#### D.1.2 SLiM settings

We simulate a diploid population of 1000 individuals, corresponding to *N*_*e*_ = 2000 chromosomes, under a Wright–Fisher model. Symmetric mutation rates, *u* = *v* = *μ*, are used at the biallelic locus, with *μ* ∈ {3 × 10^−4^, 10^−3^, 3 × 10^−3^}. In SLiM, the selection coefficient is *S* = 2*s* with a dominance coefficient of 0.5, chosen to match, to first order in selection strength, the haploid selection coefficient *s*; the parameter values are *s* ∈ {0.005,0.01,0.05}. The initial frequency is *p*(0) = 0.01, corresponding to *N*_*e*_ *p*_0_ = 20 favorable allele copies; in the weakest-selection, weakest-mutation case (*s* = 0.005, *μ* = 3 × 10^−4^), genetic drift has a relatively strong effect over 2000 generations starting from *p*(0) = 0.01. Additional runs start at the high initial frequency specified in Section 7.1, so that the experiments test convergence toward equilibrium from both sides (*p*(0) = 0.99 and *p*(0) = 0.01).

#### D.1.3. Parameter sweep

The *γ* sweep in Fig. 2C gives the following steady-state values: max|*g* − *ρ*| = 2.92 × 10^−1^ at *γ* = 0.01 (with *g* → 0.646, *ρ* → 0.938), 6.01 × 10^−2^ at *γ* = 0.1, 6.69 × 10^−3^ at *γ* = 1, 6.76 × 10^−4^ at *γ* = 10, and 6.77 × 10^−5^ at *γ* = 100. At steady state, 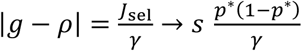, where the first relation is exact and the second holds in the strong-coupling limit; both are consistent with the numerical results. Across all 11 parameter sets, the VPG analytical solutions agree with the mean final frequencies from SLiM, with a maximum deviation of < 2 SE (for the strongest-selection case *μ* = 3 × 10^−3^, *s* = 0.05).

### D.2 Supplementary material for Experiment 2: two-locus recombination–selection dynamics

#### D.2.1 Numerical solution

The time step is *dt* = 0.01, and the simulations are run for *T* = 2000 generations. DVPG is integrated to 4000 in model time and plotted against the rescaled time *τ* = *t*/2, corresponding to 2000 generations. The recombination flux is implemented using the projection form ***J***_rec_ = *r*(*Π****g*** − ***g***). Here, *Π* denotes the projection onto the product of the one-locus marginal distributions; the maximum numerical difference between the two recombination-flux implementations is below 10^−16^. The DVPG coupling flux is treated as in Experiment 1(operator splitting with a matrix-exponential step). For *L* = 2, as a numerical cross-check, the full right-hand side (RHS) is also integrated using SciPy’s BDF solver, with no excursion outside the probability simplex and consistent results. In the absence of mutation, recombination drives the system toward linkage equilibrium: after fixation at the selected locus (*p*_2_ → 1), *D* = *p*_11_ − *p*_1_*p*_2_ → 0, where *p*_11_ is the frequency of haplotype (1,1), and *p*_1_ and *p*_2_ are the corresponding marginal allele frequencies at the two loci; this condition is used as the convergence criterion. The coupling strength is *γ* = 100.

#### D.2.2 SLiM settings

We simulate a diploid population of 1000 individuals (*N*_*e*_ = 2000 chromosomes). Mutation is switched off (*u* = *v* = 0). A single recombination interval is used, with recombination rate *r* ∈ {0.001,0.01}. Selection acts only on the second locus (simulation locus 1), with selection coefficient *s* ∈ {0.005,0.05}. In SLiM, the corresponding dominance coefficient is 0.5 (*S* = 2*s*), which matches the haploid selection coefficient to first order in selection strength. The initial haplotype frequencies are *p*_00_ = 0.45, *p*_11_ = 0.45, *p*_01_ = *p*_10_ = 0.05, corresponding to counts of [900,100,100,900] and an initial linkage disequilibrium of *D*(0) = 0.20.

#### D.2.3. Parameter sweep

The final-state *p*_11_ is evaluated over the 2 × 2 parameter grid (*s* ∈ {0.005,0.05} × *r* ∈ {0.001,0.01}): VPG predicts [0.811,0.889; 0.591,0.811], whereas the SLiM simulations yield [0.815 ± 0.031,0.883 ± 0.027; 0.547 ± 0.038,0.780 ± 0.033] across 100 independent replicates. All final-state differences are within 2 SE, consistent with finite-population sampling variability under the tested settings (matrix rows: the two recombination rates, 0.001 and 0.01; matrix columns: the two selection coefficients, 0.005 and 0.05).

### D.3 Supplementary material for Experiment 3: three-locus integrated system and stochastic extension

#### D.3.1 Numerical solution

The time step is *dt* = 0.01, and the simulations are run for *T* = 2000 generations. DVPG is integrated to 4000 in model time and plotted against the rescaled time *τ* = *t*/2, corresponding to 2000 generations. The mutation generator is the ***M*** (8 × 8) rate matrix, the recombination flux is implemented in the projection form ***J***_rec_ = *r*_1_(*Π*_0|12_***g*** − ***g***) + *r*_2_(*Π*_01|2_***g*** −***g***), and the selection flux is *J*_sel,*h*_ = *ρ*_*h*_(*U*_*h*_ − *Ū*) with the additive selection potential *U*_*h*_ = ∑_*l*_*s*_*l*_ *b*_*h,l*_. For DVPG, operator splitting with a matrix-exponential step is used for the coupling subsystem. Note that for *L* = 3 the DVPG contains a bilinear recombination projection, and the internal trial steps of the SciPy BDF solver can leave the probability simplex and cause numerical divergence; an explicit scheme with per-step clipping followed by normalization is therefore used.

#### D.3.2 SLiM settings

By default, we simulate a diploid population of 1000 individuals (with two additional populations of 500 and 3000 individuals used in the stochastic extension), corresponding to *N*_*e*_ = 2000 chromosomes. Symmetric bidirectional mutation rates *u* = *v* = 0.001 act at all three loci. Selection is locus-specific with additive effects *s* = [+0.005,0, −0.005] (locus 0 beneficial, locus 1 neutral, locus 2 deleterious); in SLiM the per-copy selection coefficients are *S* = 2*s* with a dominance coefficient of 0.5, chosen to match the haploid VPG selection coefficients to first order in selection strength. Two recombination intervals are used, with rates *r*_12_ = 0.01, *r*_23_ = 0.05. The initial haplotype frequencies are *p*_000_ = *p*_111_ = 0.4, and each of the remaining six haplotypes has frequency approximately 0.033 (0.2/6), corresponding to counts of 800, 67, and 66 haplotypes out of *N*_*e*_ = 2000 for the default population of 1000 individuals; for the 500- and 3000-individual populations the corresponding initial counts are (400, 34, 33, 33, 33, 33, 34, 400) and (2400, 200, 200, 200, 200, 200, 200, 2400), respectively. This initial configuration carries strong linkage disequilibrium across the three loci. Haplotype frequencies are recorded every 10 generations over *T* = 2000 generations.

#### D.3.3 Stochastic simulation

For the stochastic extension, the SVPG dynamics with genetic drift (Langevin noise) are integrated with an explicit Euler–Maruyama scheme with time step *Δt* = 0.01 for *T* = 2000 generations, using the same mutation, selection, and recombination parameters as the deterministic experiments. The haploid VPG effective population size (*N*_*e*_ = 1000/2000/6000 chromosomes) is set to twice the SLiM diploid population size (500, 1000 and 3000 individuals, respectively). The noise covariance is the Fisher–Rao matrix ***M***_***FR***_(***ρ***) = *diag*(***ρ***) −***ρρ***^T^ divided by the effective population size. After each integration step, negative components are clipped and the state is renormalized to the probability simplex. For each population size, 100 independent stochastic trajectories are simulated with consecutive NumPy default_rng seeds. Mean trajectories and standard errors are computed across the independent replicates and compared with the corresponding SLiM forward simulations (100 replicates).

## Code Availability

Scripts for reproducing the numerical experiments presented in this paper are available on GitHub at https://github.com/cailigd/vpg_paper.

## Acknowledgments

We thank Dr. Jiajun Zhang and Dr. Xiaochen Yu for helpful discussions and insightful comments on the manuscript. This work was supported by the National Natural Science Foundation of China (32470690).

